# Spatial polarization of endothelial ICAM-1 governs T-cell exclusion in melanoma

**DOI:** 10.64898/2026.03.19.712709

**Authors:** Ha-Ram Park, Shawn J. Kim, Nika P. Kozlov, Somnath Tagore, Lawrence W. Wu, Benjamin Izar, Minah Kim

## Abstract

An immunosuppressive tumor microenvironment limits therapeutic efficacy and worsens prognosis in melanoma. Beyond T-cell abundance and function, effective tumor control also depends on whether T cells can access malignant cells within the tumor. Although emerging evidence supports that tumor vasculature facilitates immune evasion, the vascular mechanisms that govern intratumoral T-cell positioning remain poorly defined. Using RNA sequencing of endothelial cells isolated from tumor cores versus peripheries in a mouse melanoma model, we identified intercellular adhesion molecule 1 (ICAM-1) as a candidate regulator of T-cell localization. During tumor growth, T cells shifted from a balanced core–margin distribution to marked exclusion from the core, most prominently in T cell-inflamed tumors. This spatial redistribution —less evident in other immune subsets—coincided with high expression of lymphocyte function-associated antigen-1 (LFA-1) on T cells. In parallel, endothelial ICAM-1 became enriched at the tumor periphery, where vascular integrity was compromised, as evidenced by increased vascular leakage and reduced pericyte coverage. Functionally, ICAM-1 blockade restored intratumoral T-cell infiltration, enhanced effector activity, and significantly delayed the growth of immunogenic tumors. Moreover, ICAM-1 inhibition sensitized an immune-refractory tumor to anti-PD-1 checkpoint blockade. Together, these findings identify endothelial ICAM-1 as a vascular determinant of intratumoral T-cell positioning and highlight the ICAM-1/LFA-1 axis as a modifiable checkpoint to reverse T-cell retention at the tumor periphery, thereby enhancing antitumor immunity and immunotherapy efficacy.

## Introduction

The clinical efficacy of immunotherapy is often limited by the immunosuppressive tumor microenvironment (TME)^1,2^. Although the presence of tumor-infiltrating cytotoxic T cells is a valuable prognostic and predictive biomarker, recent studies show that their spatial proximity to tumor cells—along with their overall abundance—strongly correlates with responses to immune checkpoint blockade (e.g., anti-PD-1 and anti-CTLA-4)^3–5^. Based on spatial immune profiles, tumors can be classified as inflamed, immune-excluded, or immune-desert. Inflamed tumors harbor abundant cytotoxic T cells intermingled with tumor cells throughout the parenchyma, whereas immune-desert tumors exhibit a paucity of effector T cells in both the tumor core and the invasive margin. The immune-excluded state—where T cells accumulate at the invasive margin but fail to penetrate the tumor core—is also associated with ineffective tumor control^6,7^. We recently demonstrated that disrupted vascular integrity at the tumor periphery contributes to T-cell exclusion in melanoma^8^, implicating tumor vasculature as a tractable target to promote balanced intratumoral T-cell positioning. However, the vascular mechanisms that regulate optimal T-cell distribution within tumors remain poorly defined.

Emerging evidence highlights context-dependent roles of intercellular adhesion molecule 1 (ICAM-1) within the TME in both endothelial and tumor compartments. On tumor cells, ICAM-1 has been reported to either promote metastatic dissemination or restrain tumor growth through distinct mechanisms, underscoring its bidirectional influence on tumor control^9–12^. Endothelial ICAM-1 classically mediates leukocyte adhesion and transendothelial migration; beyond facilitating entry, it can also shape T-cell effector function within the TME^13,14^. Emerging evidence indicates that heterogeneous endothelial ICAM-1 expression generates discrete transmigration hotspots that regulate leukocyte entry while preserving vascular integrity^15^. Such spatial regulation of endothelial transmigration suggests that ICAM-1 may influence not only the T-cell abundance but also the regional distribution of T cells within tumors. Whether endothelial ICAM-1 drives the emergence of T-cell exclusion during disease progression—and whether targeting ICAM-1 can reverse exclusion and enhance immunotherapy—remains unresolved.

Here, we identify endothelial ICAM-1 as a key mediator of intratumoral T-cell positioning in melanoma. As tumors grew, T cells shifted from a balanced core-margin distribution to marked exclusion from the core, most prominently in immunogenic tumors. This redistribution coincided with high expression of lymphocyte function–associated antigen-1 (LFA-1) on T cells and increased ICAM-1 on compromised peripheral tumor vessels (< 500 µm from the invasive margin). Functionally, ICAM-1 blockade restored a more balanced T-cell distribution, enhanced T-cell function, and delayed tumor growth. Moreover, ICAM-1 inhibition sensitized an immune-refractory tumor to checkpoint blockade, accompanied by improved CD8^+^ T-cell penetration toward the tumor core and increased effector activity. These findings establish endothelial ICAM-1 as a vascular determinant of T-cell localization and provide a rationale for targeting the ICAM-1/LFA-1 axis to enhance T-cell access to malignant cells and potentiate antitumor immunity.

## Materials and Methods

### Patient samples

Archived FFPE primary and metastatic melanoma tissues were obtained from the Herbert Irving Comprehensive Cancer Center (HICCC) Tissue Bank under an approved IRB protocol. Sections (5 µm or 20 µm) were prepared by the HICCC Histology Service for immunostaining.

### Mice

Pathogen-free C57BL/6J mice were purchased from the Jackson Laboratories and maintained on standard chow under specific pathogen–free conditions. Seven- to eight-week-old mice were subcutaneously injected with 2.5 × 10^5^ YUMMER1.7 cells or 5 × 10^5^ YUMM1.7 cells into the flank. Tumors were collected at ∼100 mm^3^ (day 9) and ∼900 mm^3^ (day 20-23), or at endpoint after treatment. For cell inoculation and perfusion, mice were anesthetized with intraperitoneal injections of ketamine (90–100 mg/kg) and xylazine (10–17 mg/kg). All animal experiments were approved by the Institutional Animal Care and Use Committee (IACUC) of Columbia University Irving Medical Center (CUIMC) and conducted in accordance with institutional and NIH guidelines.

### Cell culture

YUMM1.7 and YUMMER1.7 (Yale University Mouse Melanoma Exposed to Radiation), which harbor driver mutations common in melanoma (*Braf*^V600E^, *Pten*^-/-^, and *Cdkn2a*^-/-^), were originally generated by the Bosenberg laboratory at Yale University^16^ and kindly provided by Dr. Amanda Lund (NYU Langone Health)^8^. Cells were maintained in DMEM-F12 supplemented with 10% fetal bovine serum (FBS) at 37°C and 5% CO_2_. Cultures were verified as Mycoplasma-negative and used at <10 passages for all experiments.

### Treatment

Tumor-bearing mice were randomized to treatment or control groups when tumors reached ∼70 mm³ (mean ∼day 8 post-inoculation). Mice received intraperitoneal injections of anti-mouse ICAM-1 (CD54) monoclonal antibody (BioXcell, BE0020, clone YN1/1.7.4; 2.5 mg/kg), anti-PD-1 (BioXcell, BP0146, clone RMP1-14; 5 mg/kg), or rat IgG2b isotype control (BioXcell, BE0090) every other day for a total of seven doses. Tumor burden was assessed twice weekly by caliper, and volume was calculated as (length × width²)/2. Tissues were collected at terminal perfusion, 3–4 weeks post-inoculation, or humane endpoint. Humane endpoints included any of the following: tumor volume ≥ 2,000 mm³; ulceration or necrosis at the tumor site. Upon meeting endpoint criteria, treatment was discontinued, and animals were euthanized under deep anesthesia followed by cardiac perfusion with physiological buffer for downstream analyses. All procedures were conducted under an IACUC-approved protocol and in accordance with institutional guidelines.

### Mouse tumor tissue preparation

Mice were perfused through the left ventricle into the aorta with 10–20 mL of cold PBS or 1% paraformaldehyde (PFA). Tumors were processed according to the following applications. For immunofluorescence staining, tissues were fixed with 1% PFA for 1 hour on ice, incubated overnight in 30% sucrose at 4°C, embedded in OCT, and sectioned at 50 µm. Frozen blocks and sections were stored at -80°C (long-term) or -20°C (short-term). For immunohistochemistry and spatial transcriptomics, tissues were fixed in 4% PFA overnight at 4°C, dehydrated in 70% ethanol, and paraffin-embedded by the Histology Service at the HICCC. Sections were prepared at a 5–10 µm thickness.

### Immunostaining

Tumor sections (50 µm) were rinsed in PBS with 0.3% Triton X-100 (PBST), blocked with 5% donkey or goat serum (Jackson ImmunoResearch) for 1 h at room temperature, and incubated with primary antibodies overnight at 4°C. Alexa-labeled secondary antibodies were applied in PBST for 4 h at room temperature, followed by DAPI (1 mg/ml; Sigma-Aldrich, D9542) nuclear staining. Slides were mounted with Fluoromount-G (Invitrogen, 00-4958-02). Antibodies are listed in Supplementary Table S1. FFPE mouse or human melanoma sections (5 µm or 20 µm) were deparaffinized in xylene, rehydrated through a graded ethanol series (100%, 95%, 90%, 80%, 75%), and subjected to antigen retrieval in citrate buffer (40 min, steamer). Sections were blocked with endogenous peroxidase blocker and 2.5% horse or goat serum, then incubated with primary antibodies overnight at 4°C or for 3 h at room temperature. ImmPRESS® HRP Horse Anti-Rabbit IgG PLUS Polymer Kits (Vector Laboratories, MP-7801) were used for detection, and chromogenic signal was developed with DAB, followed by hematoxylin counterstaining (Sigma, GHS316). Antibodies are listed in Supplementary Table S1. Images of immunofluorescence-stained tissues were acquired using Axio Observer 7 with Apotome2 [Zeiss, 10× or 20× objective, bin 1 or 2], and analyzed with Image J v1.54. Immunohistochemistry-stained slides were scanned using Leica AT2 scanner and analyzed with QuPath v0.6.0.

### Flow cytometry

Tumors were dissected into core and peripheral regions or analyzed as whole tumors, depending on the experiment. Tissues were enzymatically digested into single-cell suspensions^8^ and analyzed on a Novocyte Penteon (Agilent). Data were processed using FlowJo v10.10.0. Antibodies are listed in Supplementary Table S1.

### Endothelial cell sorting and Bulk RNA-seq

Large tumors (∼900 mm³) were dissected into core and periphery and digested in buffer containing 2% FBS, 0.1% collagenase IV (Worthington, LS004188), and 10 U/ml DNase I. Minced tissue was filtered through 70 µm strainers and washed with MACS buffer (0.5% BSA in PBS). Endothelial cells (CD45⁻CD31⁺) were enriched by sequential magnetic cell separation (CD45-depletion, CD31-positive selection; Miltenyi Biotec), followed by fluorescence-activated cell sorting (FACS) to > 95% purity. RNA was extracted using the miRNeasy Micro Kit (Qiagen, 217084), and quality confirmed (RIN > 7.0; Agilent Bioanalyzer). Libraries were prepared using the SMARTer Stranded Total RNA-seq Kit v2 (Clontech) and sequenced on an Illumina NovaSeq 6000 at the HICCC Genome Center.

### Spatial transcriptomics analysis

Spatial transcriptomics was performed on mouse melanoma samples using the Xenium platform (10x Genomics). To maximize sample size, 1–2 mm tissue cores were collected from FFPE YUMMER1.7 tumors (11 small, ∼100 mm³; 11 large, ∼900 mm³) to construct tissue microarrays (TMA). Peripheral regions (< 500 µm from the tumor margin) and core regions (> 500 µm from the tumor margin) were sampled, yielding a total of 36 cores embedded into two TMA blocks. Sections (5 µm) were processed on Xenium slides according to the manufacturer’s protocols. Imaging and data acquisition were performed using a Xenium Analyzer by the HICCC Genome Engineering Core. A custom 407-gene panel enriched for immune and angiogenic pathways was designed based on a panel developed by Dr. Thorsten R. Mempel’s group.

### Bioinformatic analysis

#### Bulk RNA-seq

FASTQ files were quality-checked with FASTQC v0.11.9. Transcript quantification was performed using Salmon v1.4.0 (selective alignment) against the mouse GENCODE release M30 (GRCm39). Differential expression analysis was conducted with DESeq2 v1.38.3; genes were considered significant at false discovery rate (FDR) < 0.05 and |log₂ fold-change| > 0.5.

#### Xenium spatial transcriptomics

Datasets were visualized with Xenium Explorer 3. Peripheral (< 500 µm) and core regions were manually annotated and merged. Computational analyses were performed in R using Seurat v5.3.0 for dimension reduction, clustering, UMAP, and spatial transcriptomic visualization.

#### Human melanoma single-cell RNA sequencing

Single-cell RNA sequencing (scRNA-seq) data from a previously generated human melanoma cohort were analyzed at the tumor level to evaluate associations between endothelial ICAM1 expression and immune composition. To avoid pseudo-replication arising from cell-level statistical testing, each tumor was treated as an independent biological replicate, and all summary metrics were computed per tumor before statistical analyses. Endothelial cells were identified using the curated cell-type annotations provided with the dataset. Normalized gene expression values were extracted from the RNA assay, and ICAM1 expression was quantified within endothelial cells for each tumor. Two endothelial ICAM-1 metrics were evaluated: (i) the fraction of ICAM1⁺ endothelial cells, defined as the proportion of endothelial cells with detectable ICAM1 expression (normalized expression > 0), and (ii) the mean normalized ICAM-1 expression across all endothelial cells within a tumor. The fraction of ICAM-1⁺ endothelial cells was used as the primary metric in correlation analyses because it captures the extent of vascular activation across the endothelial compartment rather than expression intensity in a subset of cells. Tumor-level immune composition was quantified as the fraction of each immune lineage among all captured cells within each tumor. Total T/NK abundance was defined as the number of annotated T/NK cells divided by the total number of cells sequenced in that tumor. For fine-state analyses, annotated T/NK subsets (including CD4⁺ T cells, CD8⁺ T cells, regulatory T cells, NK cells, and additional naive or memory states) were quantified similarly at the tumor level. For each immune subset, the tumor-level fraction was used in downstream correlation testing. T-cell receptor (TCR) clonality was calculated at the tumor level using normalized Shannon entropy. For each tumor, clonotype frequencies were derived from annotated TCR sequences, and Shannon entropy (H) was computed based on the distribution of clonotype frequencies. To account for differences in clonotype counts across tumors, entropy was normalized to the theoretical maximum (log N, where N is the number of unique clonotypes), and clonality was defined as 1 − (H / log N). This normalization ensures comparability across tumors with different T-cell counts or sequencing depth. Associations between endothelial ICAM-1 metrics and tumor-level immune abundance were assessed using two-sided Spearman rank correlation tests, given the modest sample size and non-normal distribution of immune fractions. For correlation analyses involving multiple T/NK fine states, *p*-values were adjusted using the Benjamini–Hochberg FDR procedure to control for multiple hypothesis testing. Statistical significance was defined as FDR < 0.05. For clonality comparisons, tumors were stratified into ICAM1-high and ICAM1-low groups using the upper and lower quartiles of endothelial ICAM-1 expression, and differences between groups were assessed using a two-sided Wilcoxon rank-sum test. All analyses were performed in R.

### Statistical analysis

Data are presented as mean ± SEM. Unless otherwise specified, all tests were two-sided, and *P* < 0.05 was considered statistically significant. Paired two-tailed Student *t*-test was used for comparisons between regions within the same samples; unpaired two-tailed Student *t*-test was used for comparisons between independent groups. Tumor growth curves were analyzed by two-way ANOVA. Exact *p*-values are presented in each figure. Analyses were performed using GraphPad Prism v10.2.1.

## Data availability

All data supporting the findings of this study are available upon request from the corresponding author. Raw and processed mouse RNA-seq and spatial transcriptomics data generated in this study are available in Gene Expression Omnibus (GEO) under accession numbers GSE319704 and GSE319706, respectively. The human scRNA-seq dataset analyzed in this study was generated by the Benjamin Izar lab and is currently under review for publication. The data is available upon reasonable request.

## Authors’ Contributions

H.R. Park: Conceptualization, data curation, formal analysis, validation, writing—review and editing. S.J. Kim: Investigation, validation, writing—review and editing. N.P. Kozlov: Investigation, validation. S. Tagore: Patient scRNA-seq data analysis. L.W. Wu: Clinical data acquisition and analysis. B. Izar: Resources, supervision. M. Kim: Conceptualization, data curation, formal analysis, investigation, supervision, funding acquisition, writing—original draft, writing—review and editing.

## Acknowledgement

We thank Amanda Lund (NYU Langone Health) for providing YUMMER1.7 and YUMM1.7 cells and Thorsten R. Mempel and Lukas M. Altenburger (Massachusetts General Hospital) for sharing the customized Xenium gene panel. We are grateful to Michael Kissner, Director of Operations of the Columbia Stem Cell Initiative Flow Cytometry Core Facility, for assistance and technical advice with flow cytometry; the DataBase Shared Resource at HICCC for screening patient melanoma tissue samples; the Molecular Pathology Shared Resource at HICCC for providing human melanoma tissues and pathological analyses; and the Single Cell Analysis Core at HICCC for support with RNA-seq experiments in melanoma models. This work was supported by grants from the National Cancer Institute (R37 CA266270 and R21 CA293766 to M.K.), the Melanoma Research Alliance (1267014 to M.K.), and the American Cancer Society (RSG-22-167-01-MM to M.K.).

## Results

### Spatial distribution of CD8^+^ T cells and its evolution during melanoma progression

We first profiled pretreatment human melanomas (n = 37; primary and metastatic) to define CD8^+^ T-cell spatial patterns within the TME. Based on CD8^+^ T-cell localization, tumors were classified into three phenotypes (Supplementary Table S2): infiltrated (relatively even distribution across periphery and core; n = 14, ∼38%), excluded (peripheral accumulation with relative absence from the core; n = 8, ∼22%), and desert (minimal infiltration throughout; n = 15, ∼41%) (**Fig. 1A** and **B**). Among melanoma patients treated with immune checkpoint inhibitor (ICI; n = 20; anti-PD-1 or anti-CTLA-4), response rates followed a graded pattern: highest in infiltrated tumors (75%), intermediate in excluded tumors (50%), and lowest in desert tumors (17%) (**Fig. 1C**). The excluded group showed a trend toward poorer overall survival compared with the infiltrated group (median OS 5.6 years vs. not reached; hazard ratio = 2.54), although this difference did not reach statistical significance (Supplementary Fig. S1).

**Figure 1.**
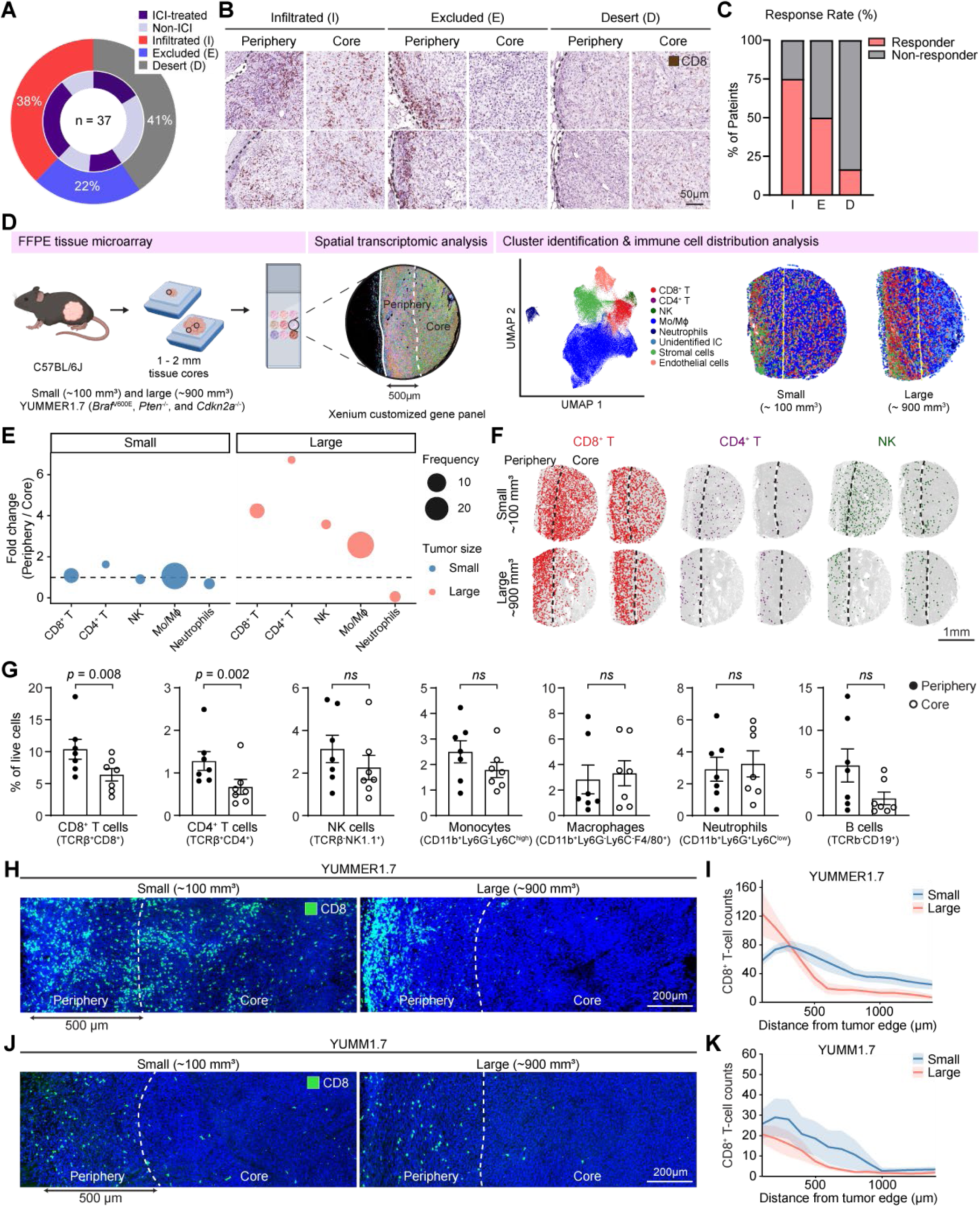
Spatial distribution of CD8^+^ T cells in human and murine melanoma and its evolution during tumor progression. **(A)** Two-ring donut chart summarizing spatial phenotypes and treatment status in 37 primary/metastatic melanomas from patients (see Supplementary Table S2). Outer ring: Infiltrated, Excluded, Desert; inner ring: ICI-treated vs non-ICI. **(B)** Representative CD8 immunohistochemistry from periphery and core illustrating the three spatial phenotypes in human melanoma. Scale bar, 50 µm. **(C)** Proportion of ICI responders and non-responders within each spatial phenotype among patients with available response data (n = 20). **(D)** Workflow: C57BL/6J mice were subcutaneously inoculated with 2.5 × 10⁵ YUMMER1.7 cells; tumors were harvested at ∼100 mm³ (small, day 9) and ∼900 mm³ (large, days 20–23). FFPE tissue microarrays were generated from 1–2 mm cores sampled from tumor periphery and core, followed by Xenium in situ analysis, cluster identification (UMAP), and spatial mapping. Examples shown for small and large tumors. Mo/Mϕ, monocytes/macrophages. **(E)** Fold change of immune populations (CD8^+^ T, CD4^+^ T, NK, Mo/Mϕ, and neutrophils) in the periphery relative to the core, comparing small (blue) and large tumors (red). Fold change was calculated as the ratio of the frequency of each cell population in the periphery to that in the core. A value of 1 (dashed line) indicates equal distribution. Dot size represents the overall frequency of each cell population across all tumors. **(F)** Representative spatial maps of CD8^+^ T, CD4^+^ T, and NK in the periphery and core of small and large tumors. Dashed lines mark the periphery–core boundary (500 µm from the tumor margin). Scale bar, 1 mm. **(G)** Flow cytometric comparison of immune-cell frequencies in periphery vs core of large YUMMER1.7 tumors (n = 7), including CD8^+^ T cells (TCRβ^+^CD8^+^), CD4^+^ T cells (TCRβ^+^CD4^+^), NK cells (TCRβ^-^NK1.1^+^), monocytes (CD11b^+^Ly6G^-^Ly6C^high^), macrophages(CD11b^+^Ly6G^-^Ly6C^-^F4/80^+^), neutrophils (CD11b^+^Ly6G^+^Ly6C^low^), and B cells (TCRβ^-^CD19^+^) among live cells. Each point represents one mouse; bars show mean ± SEM; *p*-values shown; ns, not significant. Statistical analysis was performed using a paired two-tailed Student’s t-test. **(H-K)** Representative CD8 immunofluorescence staining of small and large YUMMER1.7 (H,I) and YUMM1.7 (J,K). Dashed lines indicate the periphery–core boundary (H,J). Corresponding quantification of CD8^+^ T-cell density (cells per 50 mm^2^) as a function of distance from the tumor margin is shown for YUMMER1.7 (n = 7 – 8) and YUMM1.7 (n = 5 – 6) tumors (I, K). Scale bar, 200 µm.

To determine how these spatial immune profiles change with tumor progression, we mapped immune-cell distributions in the syngeneic YUMMER1.7 mouse melanoma model. YUMMER1.7 tumors harbor *Braf*^V600E^*, Pten*^-/-^, and *Cdkn2a*^-/-^ alterations and exhibit a T cell-inflamed phenotype driven by UV-induced somatic mutations^16^. Our previous study showed that despite the abundance of CD8^+^ T cells, YUMMER1.7 frequently exhibits immune exclusion, with CD8^+^ T cells retained at the tumor periphery^8^. Tumors were harvested at approximately 100 mm³ (“small”) and 900 mm³ (“large”). Tissue microarrays were constructed from 1–2 mm cores sampled at the tumor margins, including periphery (< 500 μm from the edge) and core regions; for large tumors, additional deep-core punches from the central region were included (Supplementary Fig. S2A). Spatial transcriptomics (10x Genomics Xenium) using a custom gene panel was then used to resolve immune positioning (**Fig. 1D**). Unsupervised clustering identified major lineages, including CD8^+^ T cells (Cd8a, Cd8b1, Trac), CD4^+^ T cells (Cd4, Foxp3), NK cells (Ncr1, Klrb1c, Klrc1), monocytes/macrophages (Itgam, Cd14, Cd68), neutrophils (S100a9, Cxcr2), and endothelial cells (Cdh5, Pecam1, Kdr) (Supplementary Fig. S2B). We quantified spatial enrichment as the periphery-to-core fold change in immune-cell cluster frequencies. In small tumors, immune populations were largely comparable between the periphery and core. In contrast, large tumors exhibited a marked shift toward periphery-enriched immune distributions, consistent with immune exclusion from the tumor core. Among lymphocytes, CD8^+^ T cells were the most abundant subset. CD4^+^ T cells showed the greatest periphery-to-core fold change, followed by CD8^+^ T cells and NK cells, whereas monocytes/macrophages —despite their overall abundance—displayed relatively modest spatial exclusion (**Fig. 1E**). Spatial maps of CD8^+^ T cells, CD4^+^ T cells, and NK cells reflected these trends: small tumors showed relatively uniform distributions, whereas large tumors displayed pronounced peripheral enrichment (**Fig. 1F**).

To validate these spatial patterns, we performed flow cytometry on dissected regions from large tumors (>900 mm³). CD8^+^ T-cell frequencies were significantly higher at the periphery than in the core (**Fig. 1F**). CD4^+^ T cells were less abundant but showed a similar pattern of exclusion, whereas NK cells, B cells, monocytes, macrophages, and neutrophils did not show significant periphery-to-core differences (**Fig. 1G**). Consistent with these findings, immunofluorescence staining supported a transition of CD8^+^ T cells from relatively uniform infiltration to progressive peripheral accumulation as YUMMER1.7 tumors grew, recapitulating the clinically relevant immune-excluded TME (**Fig. 1H**). Distance-based quantification from the tumor margin inward further supported a gradual emergence of core exclusion with tumor growth (**Fig. 1I**). To assess T-cell positioning in immunologically “cold” tumors, we also analyzed YUMM1.7 melanomas, which contain markedly fewer T cells, as reported previously^8,16^. As expected, T-cell frequencies were substantially lower than in YUMMER1.7 tumors. Notably, unlike YUMMER1.7, YUMM1.7 tumors showed peripheral retention of CD8^+^ T cells even at the small-tumor, and this pattern persisted as tumors enlarged (**Fig. 1J** and **K**).

### Spatial enrichment of endothelial transcripts at the tumor periphery

To identify endothelial determinants of T-cell exclusion in melanoma, we performed RNA-seq on endothelial cells (CD45^-^CD31^+^) isolated from the periphery and core of large YUMMER1.7 tumors (> 900 mm^3^), which exhibit prominent CD8^+^ T-cell accumulation at the tumor margin (**Fig. 2A**). Gene Ontology analysis (periphery vs. core) revealed significant enrichment of pathways related to cell adhesion (GO:0045785), leukocyte migration (GO:0002585), extracellular matrix organization (GO:0030198), chemotaxis (GO:0006935), inflammatory response (GO:0006954), and endothelial cell migration (GO:0043542) in the peripheral endothelium relative to the core (**Fig. 2B**). These enrichments indicate an activated and remodeled vascular state at the tumor margin that is pro-adhesive and chemotactic, consistent with preferential leukocyte tethering and retention at the periphery. Consistent with these enriched pathways, curated gene modules showed elevated peripheral expression of transcripts associated with vascular development (*Adgrg1, Aplnr, Clec14a, Foxm1*), extracellular matrix remodeling (*Fbln2*, *Col14a1, Col1a1, Ccdc80*), inflammatory programs (*Afap1l2, C1qtnf3, Lpl, Tnfsf8*), chemotaxis (*Cxcl12, Cxcl13*, *Cxcl14,* and *Epha4*), and cell adhesion/migration (e.g., *Icam1, Ackr1/DARC, Elmo1*) (**Fig. 2C**). In contrast, several transcripts were relatively enriched in the tumor core and downregulated at the periphery, including *Il33, Lgals3 (Galectin-3), Mmp1a,* and *Slpi*. A volcano plot comparing peripheral versus core endothelium (log₂FC > 0.5, FDR < 0.05) highlighted key genes upregulated at the tumor periphery (**Fig. 2D**). Among immune adhesion and trafficking molecules, *Icam1* emerged as one of the most significantly induced transcripts in peripheral endothelial cells. Because endothelial ICAM-1 is a well-established mediator of T-cell adhesion and transendothelial migration, yet its role in shaping intratumoral T-cell positioning remains unclear, we prioritized ICAM-1 for further investigation. To assess clinical relevance, we analyzed a previously generated human melanoma scRNA-seq cohort (n = 25 tumors) and found that the fraction of ICAM-1⁺ endothelial cells showed a modest positive association with overall T/NK abundance (Spearman ρ = 0.38, two-sided p = 0.058; Supplementary Fig. S3A). In contrast, endothelial ICAM-1 was not significantly associated with enrichment of specific T/NK fine states after FDR correction, nor with tumor-level TCR clonality (Supplementary Fig. S3B,C), consistent with a role in immune access rather than differentiation or antigen-driven expansion. Together, these observations motivated validation of ICAM-1 spatial expression during tumor growth and testing whether polarized endothelial ICAM-1 promotes peripheral retention of CD8⁺ T cells.

**Figure 2.**
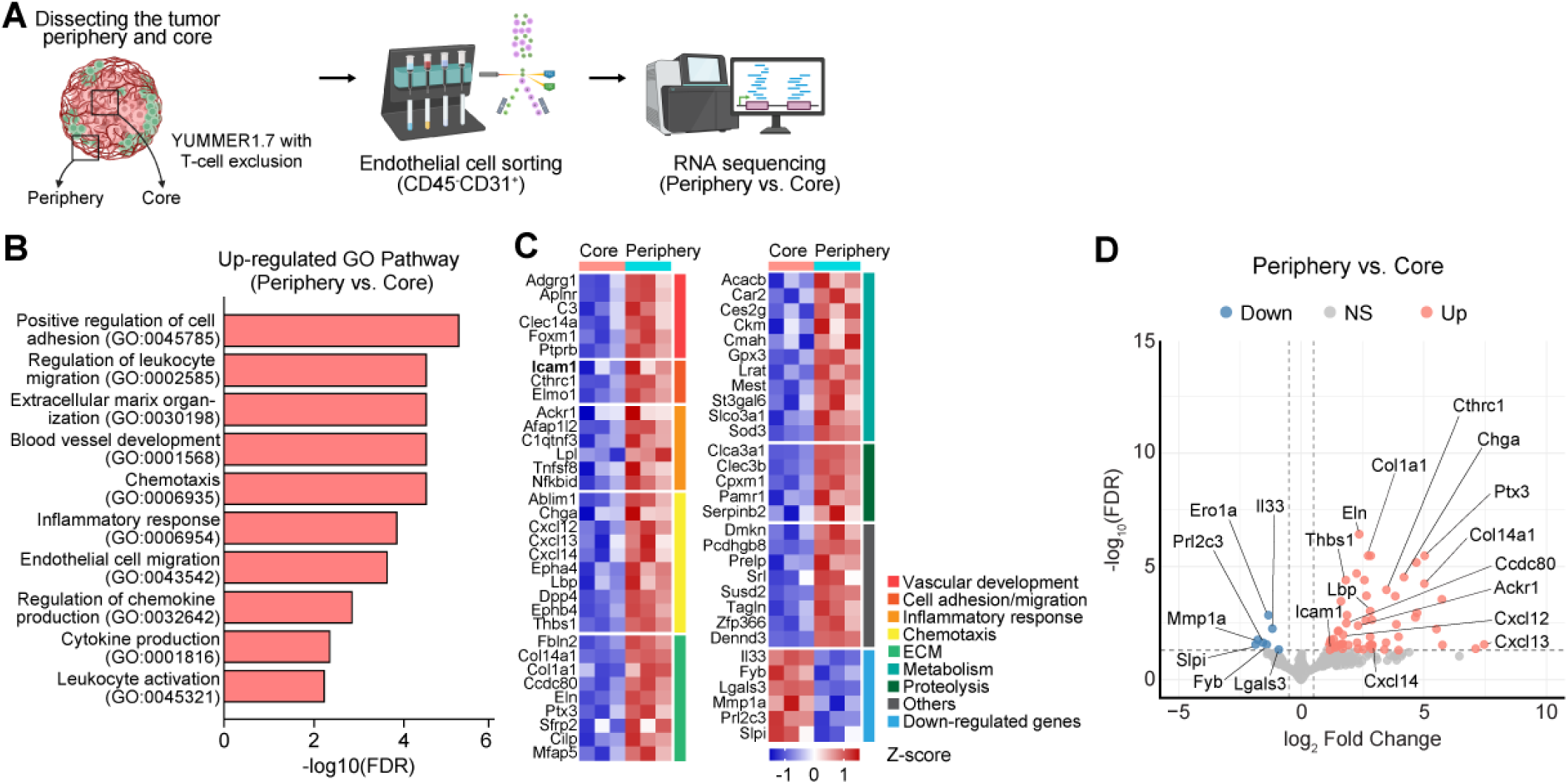
Endothelial transcripts enriched at the tumor periphery. **(A**) Workflow for endothelial cell isolation and bulk RNA sequencing. Large YUMMER1.7 melanomas (∼900 mm³) were dissected into periphery and core; endothelial cells (CD45⁻CD31⁺) were isolated from each region and profiled by bulk RNA sequencing. **(B)** Gene Ontology enrichment for Biological Process among genes upregulated in peripheral endothelial cells vs core endothelial cells. Bars show the top terms ranked by –log₁₀ false discovery rate (FDR). **(C)** Heatmap of representative differentially expressed genes, grouped by function. Columns are biological replicates from core and periphery; values are z-scored expression. **(D)** Volcano plot for periphery vs core differential expression. Points are colored up (red), down (blue), or not significant (gray). Significance was filtered by FDR < 0.05 and |log_2_ fold change| > 0.5. ECM, extracellular matrix.

### Spatial polarization of endothelial ICAM-1 expression during immunogenic tumor growth associates with vascular destabilization

To assess endothelial adhesion molecules in tumors with T-cell exclusion, we analyzed CD45^-^CD31^+^ endothelial cells from the periphery and core of YUMMER1.7 melanomas by flow cytometry. Among the adhesion molecules tested, ICAM-1 showed the highest prevalence and was significantly enriched at the tumor periphery (**Fig. 3A**), consistent with our RNA-seq results. P-selectin was also elevated at the periphery, whereas VCAM-1 and E-selectin showed no noticeable regional differences. Because LFA-1 (αLβ2) mediates post-arrest crawling and transendothelial migration^17^, we profiled its α subunit (CD11a) across intratumoral leukocytes. CD11a expression (Δ geometric MFI) was higher on CD8⁺ and CD4⁺ T cells—the subsets most strongly excluded from the core in large tumors—than on other immune populations, supporting a functional ICAM-1/LFA-1 axis relevant to intratumoral T-cell positioning (**Fig. 3B**).

**Figure 3.**
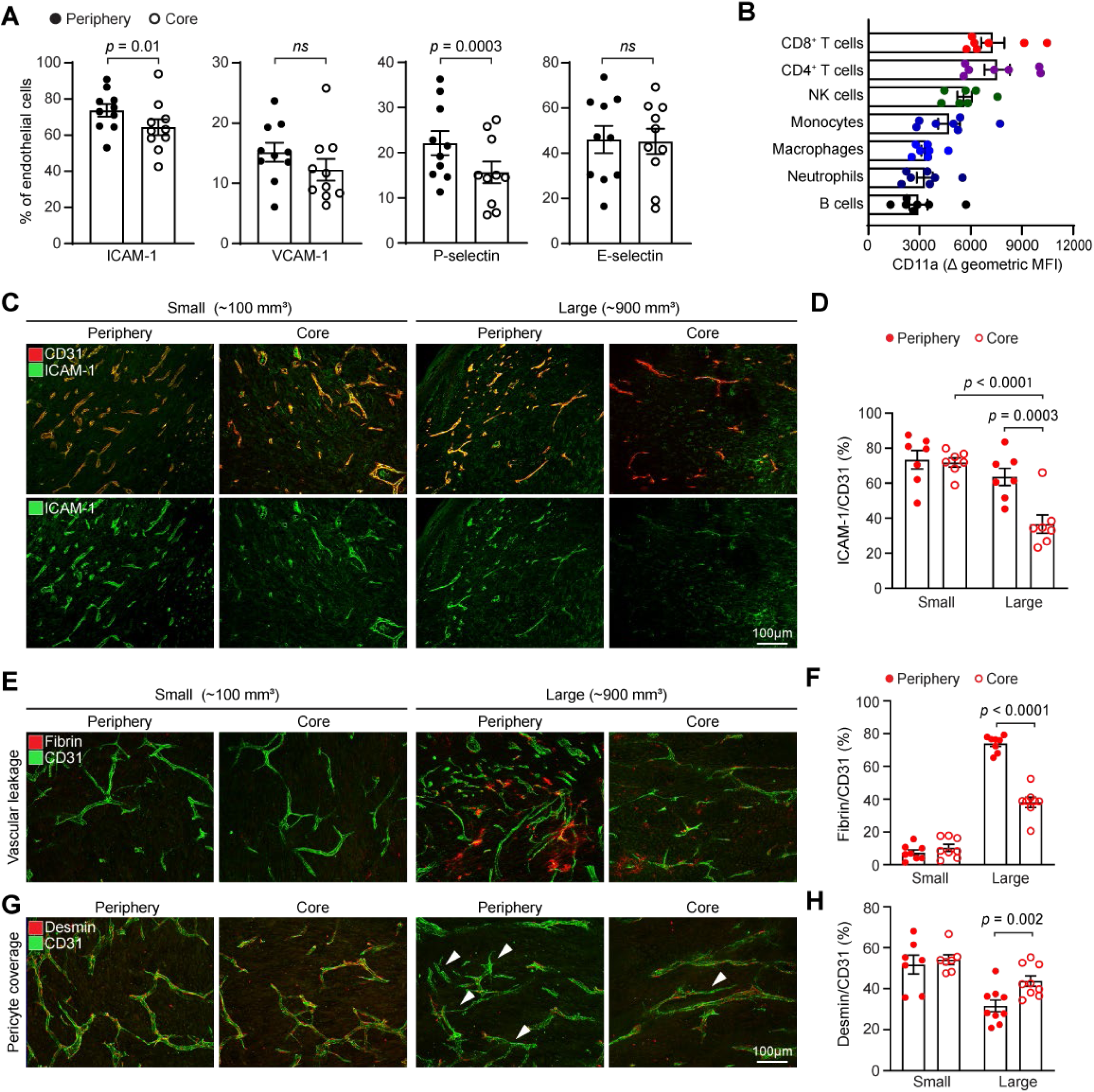
Endothelial ICAM-1 is enriched at the tumor periphery and links to vascular destabilization during immunogenic tumor growth. **(A)** Flow cytometry analysis showing frequencies of CD31⁺ endothelial cells expressing ICAM-1, VCAM-1, P-selectin, or E-selectin from the periphery vs. core of large (∼900 mm³) YUMMER1.7 tumors. Statistical analysis was performed using a paired two-tailed Student’s t-test. **(B)** Expression of CD11a (LFA-1 α-subunit; Δ geometric MFI) on the indicated tumor-infiltrating immune populations. **(C, D)** Representative immunofluorescence showing ICAM-1 (green) on CD31⁺ vessels (red) in small (∼100 mm³) and large (∼900 mm³) tumors at periphery and core (C) and corresponding quantification of ICAM-1⁺/CD31⁺ area (%) (D). Scale bar in C, 100 µm. **(E-H)** Representative images of extravasated fibrin(ogen) (red) (E) and pericyte (desmin, red) (G) relative to CD31 (green) in small and large tumors at periphery and core and corresponding quantification of fibrin/CD31 area (%) (F) and desmin/CD31 area (%) (H), respectively. Scale bar for E and G, 100 µm. Statistical analysis was performed using paired or unpaired two-tailed Student’s t-test. Each point represents one mouse; bars show ± SEM.

We then asked whether the spatial pattern of ICAM-1 changes as tumors grow. In small tumors (∼100 mm³), endothelial ICAM-1 staining was relatively uniform across the periphery and core. In large tumors (∼900 mm³), ICAM-1 became polarized to the periphery, with a marked reduction on endothelial cells in the core (**Fig. 3C**). Although ICAM-1 immunoreactivity was also detected in non-endothelial cells, the predominant signal localized to CD31^+^ endothelium. Quantification of endothelial-associated ICAM-1 confirmed a significant periphery–core difference in large tumors (*p* = 0.0003) and a pronounced decline in ICAM-1 in the core endothelium from small to large tumors (*p* < 0.0001) (**Fig. 3D**). To determine whether this polarization of ICAM-1 expression accompanies changes in vascular integrity, we assessed fibrin deposition as a readout of vascular leakiness. Small tumors exhibited minimal leakage in both regions, whereas large tumors showed pronounced peripheral leakage relative to the core (*p* < 0.0001) (**Fig. 3E** and **F**). In parallel, pericyte coverage decreased with progression and was lowest at the periphery of large tumors (**Fig. 3G** and **H**). Together, these data show that tumor growth is accompanied by vascular destabilization at the margin and elevated ICAM-1 expression, aligning with—and potentially contributing to—the peripheral retention and core exclusion of T cells.

### ICAM-1 blockade results in balanced CD8^+^ T-cell distribution and delays tumor growth

Given that CD8⁺ T-cell exclusion coincided with peripheral enrichment of endothelial ICAM-1, we next investigated whether targeting the ICAM-1/LFA-1 axis could reshape T-cell spatial organization and affect tumor progression. Mice bearing established YUMMER1.7 melanomas were treated with anti-ICAM-1 or IgG for approximately two weeks, beginning on day 8–10 after tumor implantation (**Fig. 4A**). Anti-ICAM-1 treatment significantly slowed tumor growth compared with control treatment (**Fig. 4B** and **C**). To assess intratumoral CD8⁺ T-cell distribution, we performed immunofluorescence staining for CD8 across the periphery–core axis. In control tumors, CD8⁺ T cells were predominantly confined to the periphery; in contrast, anti-ICAM-1 treatment produced a more uniform distribution (**Fig. 4D**). Quantification confirmed a pronounced periphery–core disparity in control tumors, whereas ICAM-1 blockade increased CD8⁺ T-cell counts in both the periphery and the core —with a larger increase in the core—thereby reducing regional imbalance (**Fig. 4E**). Flow cytometry corroborated these findings: the frequency of CD8⁺ T cells among live cells increased from 2.75% to 4.13% upon ICAM-1 inhibition (∼ 1.5-fold) (**Fig. 4F**). This was accompanied by an increased proportion of granzyme B⁺CD8⁺ T cells, indicating enhanced cytotoxic function (**Fig. 4G**). In contrast, the overall frequency of CD45^+^ immune cells, CD4^+^ T cells, and regulatory T cells (Foxp3^+^CD4^+^) remained unchanged. (**Fig. 4H-J**). Together, these results demonstrate that the ICAM-1/LFA-1 axis governs intratumoral CD8⁺ T-cell positioning and effector activity. Endothelial ICAM-1, therefore, acts as a vascular checkpoint that restricts CD8⁺ T-cell access to the tumor core, highlighting its therapeutic potential for overcoming immune exclusion and potentiating antitumor immunity.

**Figure 4.**
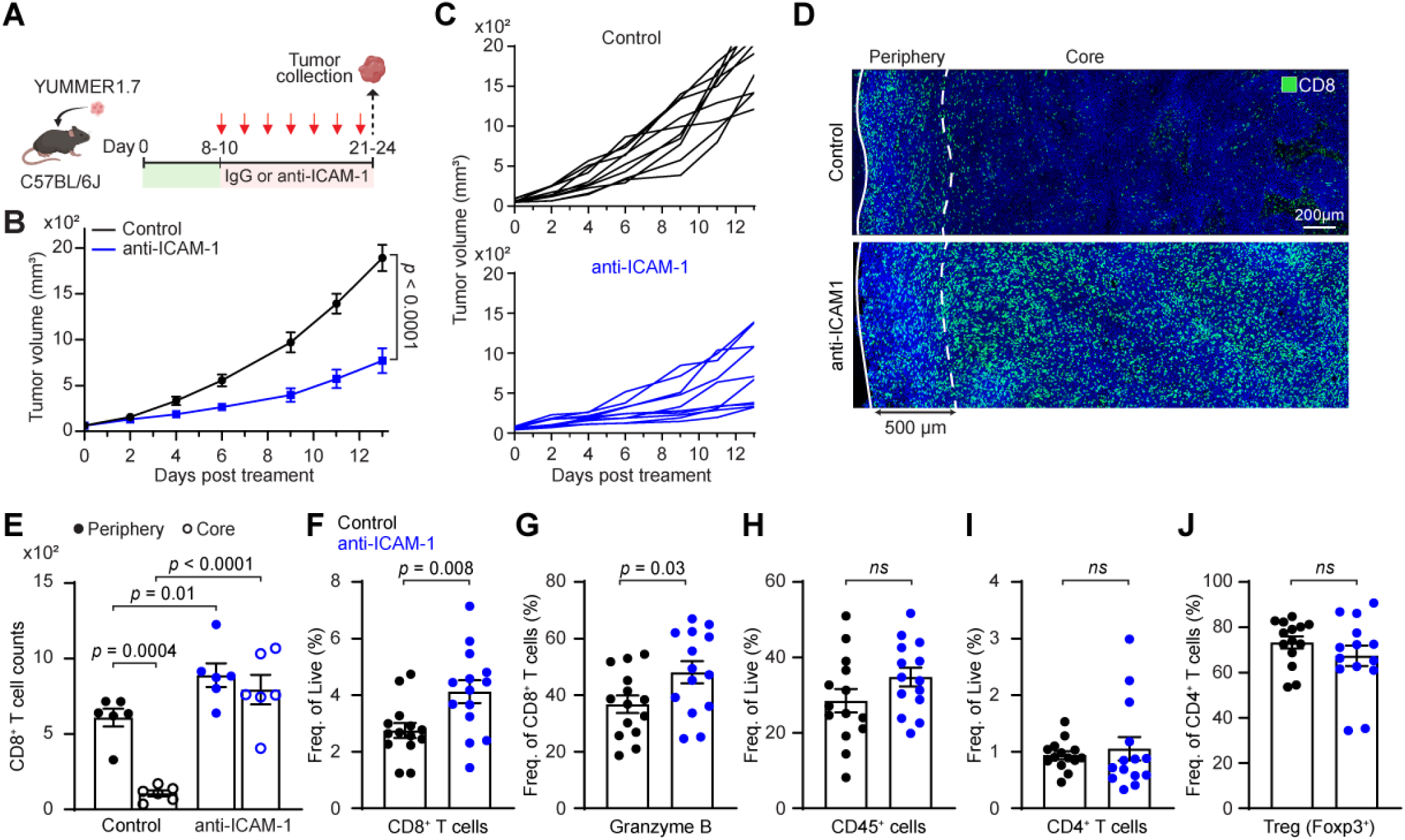
ICAM-1 blockade slows tumor growth and redistributes CD8^+^ T cells to the tumor core. **(A)** Schematic diagram of treatment. 2.5 × 10^5^ YUMMER1.7 cells were subcutaneously inoculated in C57BL/6 mice. When tumors reached ∼70 mm³ (days 8–10), mice were treated with anti-ICAM-1 or rat IgG isotype control (2.5 mg/kg, i.p.) every other day for seven doses; tumors were collected on days 21–24. **(B-C)** Tumor growth curves showing group means ± SEM (B) and individual trajectories (C) for IgG (black, n = 10) and anti-ICAM-1 (blue, n = 10) pooled from three independent experiments. *P*-value at the final time point by two-way ANOVA. **(D)** Representative immunofluorescence showing CD8⁺ T-cell distribution (green) across periphery to core in IgG- and anti-ICAM-1–treated tumors. Solid line marks the tumor boundary, and dashed line indicates the periphery–core boundary. Scale bar, 200 µm. **(E)** Quantification of CD8⁺ T-cell counts in periphery vs core (IgG, n = 6; anti-ICAM-1, n = 6). Statistical analysis was performed using unpaired or paired student two-tailed Student’s t-test. (**F-J**) Flow cytometric analysis of whole tumors comparing the frequency of CD8^+^ T cells among live cells (F), Granzyme B^+^ cells among CD8^+^ T cells (G), and CD45^+^ immune cells among live cells (H), CD4^+^ T cells (I) among live cells, and regulatory T cells (Treg; CD4^+^Foxp3^+^) among CD4^+^ T cells (J) between IgG (black, n = 14) and anti-ICAM-1 (blue, n =14). Statistical analysis was performed unpaired two-tailed Student’s t-test. Each point represents one mouse; bars show mean ± SEM.

### ICAM-1 blockade sensitizes an immune-refractory tumor to anti-PD-1 therapy

To determine whether ICAM-1 blockade can overcome resistance to immune checkpoint inhibition, we used the immune-refractory YUMM1.7 melanoma model, which exhibits sparse baseline T-cell infiltration. Anti-ICAM-1 and anti-PD-1 antibodies were administered either alone or in combination starting from day 8-10 after tumor implantation and continued for ∼2 weeks (**Fig. 5A**). Neither anti-ICAM-1 nor anti-PD-1 monotherapy delayed tumor growth; however, combined ICAM-1 and PD-1 blockade markedly slowed tumor progression compared with all other groups (**Fig. 5B**). Flow cytometry analysis confirmed that CD8^+^ T-cell infiltration in IgG-treated YUMM1.7 tumors was extremely low at baseline (0.14% of live cells). Anti-ICAM-1 or anti-PD-1 alone resulted in only modest increases in CD8^+^ T-cell frequencies (0.24% and 0.60%, respectively), whereas combination treatment increased CD8^+^ T-cells ∼8-fold to 1.18% (**Fig. 5C**). Although overall CD8^+^ T-cell abundance remained low even with combination therapy, infiltrating CD8^+^ T-cells displayed a more activated/cytotoxic phenotype, including higher proportions of granzyme B^+^ cells, increased CD62L^+^CD44^+^ effector memory cells, and increased CD69^+^CD8^+^ cells relative to the other treatment groups (**Fig. 5D-F**). Monotherapy also modestly improved these functional readouts. In contrast, the overall frequencies of CD45^+^ immune cells, CD4^+^ T cells, and regulatory T cells (Foxp3^+^CD4^+^) remained unchanged across treatment groups (Supplementary Fig. S4). Collectively, these results indicate that ICAM-1 blockade enhances both the abundance and functional state of CD8^+^ T cells in the context of PD-1 blockade, thereby sensitizing tumors resistant to anti-PD-1 therapy.

**Figure 5.**
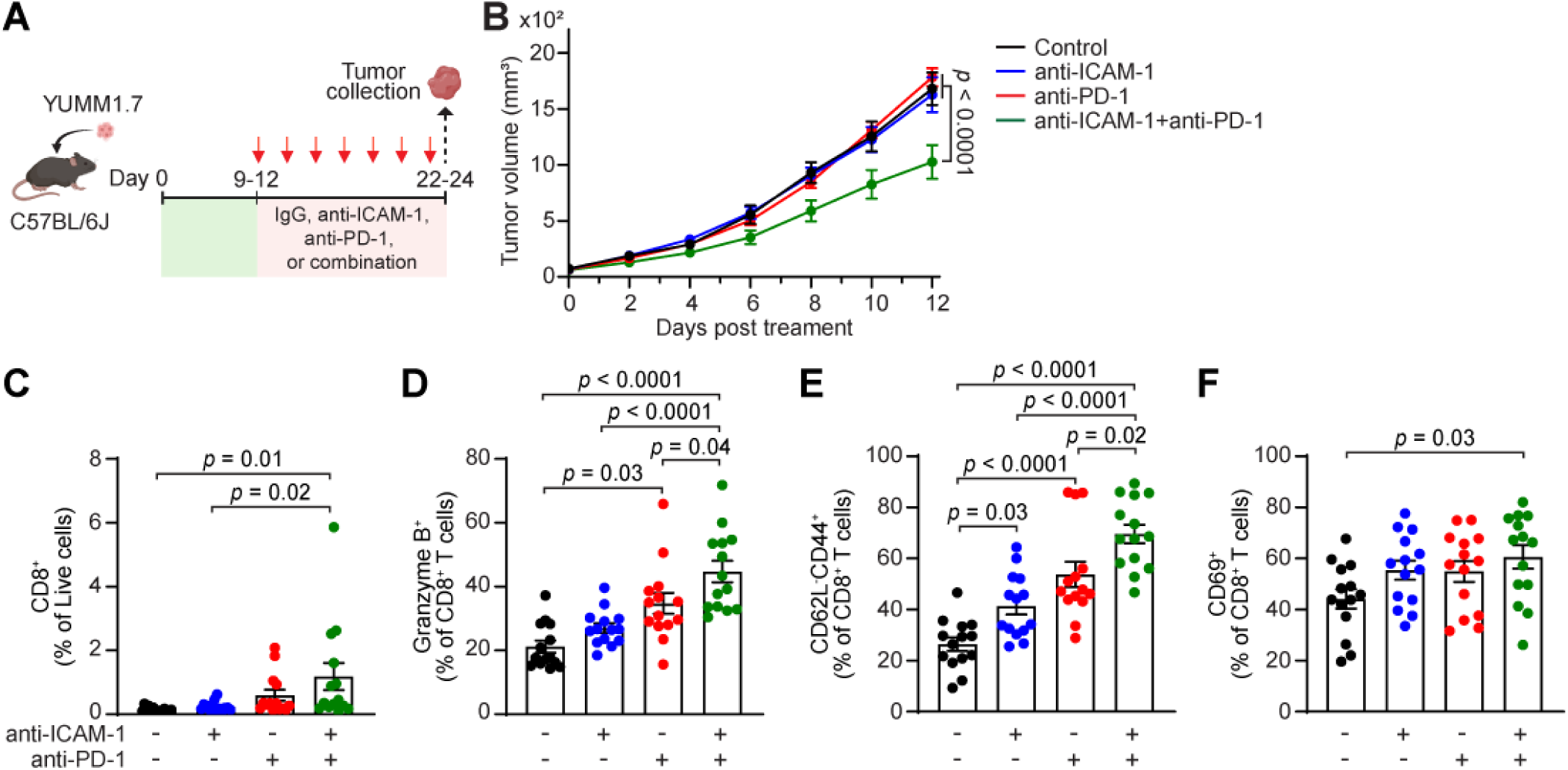
ICAM-1 blockade sensitizes tumors resistant to anti-PD-1 therapy. **(A)** Schematic diagram of treatment. 5 × 10^5^ YUMM1.7 cells were subcutaneously inoculated in C57BL/6 mice. When tumors reached ∼70mm³ (days 9–12), mice were intraperitoneally treated with rat IgG isotype control, anti-ICAM-1 (2.5 mg/kg), anti-PD-1 (5mg/kg), or combination (anti-ICAM-1 [2.5mg/kg] and anti-PD-1 [5mg/kg]) every other day for seven doses; tumors were collected on days 22-25. **(B)** Tumor growth curve showing group means ± SEM for IgG (black, n = 10), anti-ICAM-1 (blue, n = 10), anti-PD-1 (red, n = 10), and combined treatment (anti-ICAM-1 plus anti-PD-1; green, n = 10). *P*-value at the final time point by two-way ANOVA. **(C-F)** Flow cytometry analysis of whole tumors comparing frequency of CD8^+^ T cells among live cells (C) and proportions of activated CD8^+^ T cells with granzyme B^+^ (D), CD62^-^CD44^+^ (effector memory cells) (E), and CD69^+^ (F) among CD8^+^ T-cells (IgG, n = 14; anti-ICAM-1, n = 14; anti-PD-1, n =14; anti-ICAM-1 plus anti-PD-1, n = 14). For C-F, each point represents one mouse; bars show ± SEM. Statistical analysis was performed using one-way ANOVA.

## Discussion

The role of endothelial ICAM-1 in shaping the spatial organization of intratumoral T cells—particularly their exclusion from tumor cores—has remained unclear. Here, we show that endothelial ICAM-1 acts as a key regulator of T-cell positioning during melanoma progression. Endothelial RNA-seq identified peripheral enrichment of adhesion, migration, and inflammatory programs, with *Icam1* among the most periphery-enriched transcripts. In early tumors, endothelial ICAM-1 was uniformly expressed, and CD8⁺ T cells were evenly distributed across the tumor. As tumors enlarged, endothelial ICAM-1 became spatially polarized at the periphery, coinciding with CD8⁺ T-cell retention at the margin and high expression of LFA-1 (CD11a) on T cells. Functionally, ICAM-1 blockade increased intratumoral CD8⁺T-cell abundance, reduced peripheral skewing by redistributing CD8⁺ T cells toward the core, and delayed growth of immunogenic tumors. In an immune-refractory setting, ICAM-1 inhibition also sensitized an immune-refractory tumor to anti-PD-1 checkpoint blockade. Together, these data identify endothelial ICAM-1 as a vascular gatekeeper of antitumor immunity and a tractable target to overcome T-cell exclusion and enhance immunotherapy.

Our findings align with the canonical role of the LFA-1/ICAM-1 axis in T-cell arrest and diapedesis and extend this paradigm by showing that regional polarization of endothelial ICAM-1 can facilitate T-cell entry and/or retention at peripheral vessels, thereby limiting access to the core. Prior work has demonstrated that compromised vascular integrity impairs efficient T cell trafficking, whereas vascular stabilization enhances T cell infiltration and sensitizes tumors to ICI therapy. In melanoma, we previously linked angiopoietin-2-driven vascular destabilization to T-cell exclusion and immunotherapy resistance^8^. Consistent with these reports, we observed pronounced vascular leakage and reduced pericyte coverage at the periphery, where endothelial ICAM-1 and T-cell accumulation were most prominent.

These observations suggest that ICAM-1 polarization and vascular destabilization co-emerge at the tumor margin and jointly shape the spatial organization of immune cells within the TME. Further work should define upstream regulators of ICAM-1 polarization, including hypoxia, shear stress, angiopoietin/TIE2 signaling, and inflammatory cytokines such as TNF-α and IFN-γ.

ICAM-1 biology is multifaceted and highly context-dependent. On tumor cells, ICAM-1 has been reported to promote tumor progression by strengthening tumor–tumor and tumor–endothelium adhesion, thereby facilitating collective migration and metastasis —illustrating tumor-intrinsic pro-metastatic functions^9,11,22^. Conversely, tumor-cell ICAM-1 can enhance antitumor immune responses and ICI therapy by supporting the homing and activation of tumor-specific T cells^12,23,24^ and by stabilizing ICAM-1-LFA-1-mediated immune synapse formation, thereby strengthening effective T-cell-tumor engagement^25,26^. In contrast, endothelial ICAM-1 is a well-established mediator of leukocyte adhesion, transendothelial migration, and immune activation^27,28^. In melanoma, our immunofluorescent data indicate that ICAM-1 expression is predominantly endothelial, positioning it as an endothelial–immune interface rather than a tumor-intrinsic determinant. This pattern may differ in other cancers, where prevalent tumor-cell ICAM-1 may directly regulate immune recognition or metastasis. Therefore, dissecting cell–type–specific ICAM–1 activity is essential for understanding its net contribution to tumor immunity, and for guiding the selection of cancer types most likely to benefit from ICAM-1-targeting strategies. While our study employed systemic ICAM-1 blockade, future work using cell–type–specific targeting will be needed to delineate the relative contributions of endothelial versus non-endothelial compartments. Moreover, given the critical role of endothelial ICAM-1 in leukocyte trafficking, careful optimization of ICAM-1 inhibition—particularly dosing and timing—will be required to promote balanced T-cell entry across the tumor parenchyma without impeding overall immune access.

The stage-dependent redistribution of ICAM-1—uniform in early lesions but increasingly restricted to peripheral vessels in large tumors—may help explain the variable outcomes of ICAM-1–targeting strategies across tumor contexts. Our data support a model in which peripheral endothelial ICAM-1 creates dominant adhesion/retention hubs, limiting effector T-cell access to the tumor core. Consistent with this, previous work showed that ICAM-1-LFA-1-dependent CD8^+^ T-cell retention within tumors limits recirculation to draining lymph nodes^29^, supporting the idea that endothelial ICAM-1-mediated adhesion traps CD8^+^ T cells at the margin, limits their inward redistribution, and reduces tumor-cell engagement. Accordingly, ICAM-1 blockade relieves peripheral sequestration, enabling deeper CD8⁺ T-cell infiltration and enhancing effector function, thereby improving tumor control in T cell-inflamed tumors. In contrast, in immunologically “cold” YUMM1.7 tumors, ICAM-1 blockade alone was insufficient, whereas combined ICAM-1 and PD-1 inhibition was effective—suggesting that both a minimal quantitative threshold of intratumoral T cells and improved function are required for meaningful tumor control. This aligns with reports of enhanced efficacy with combined anti-ICAM-1 and ICI therapy^30,31^.

Together, these findings motivate cell type–targeted disruption of the ICAM-1/LFA-1 axis and rational combinations with ICI and/or vascular-normalizing approaches to increase intratumoral T-cell access and reinvigorate antitumor immunity. In summary, our study adds a spatial framework to ICAM-1 biology in melanoma, helps reconcile the context-dependent roles, and provides a mechanistic rationale for pairing ICAM-1/LFA-1–modulating therapies with immunotherapy or anti-angiogenic strategies to overcome immune exclusion and improve melanoma outcomes.

## Notes

### Competing Interest Statement

The authors have declared no competing interest.

