## Supplementary Materials for "Spatial polarization of endothelial ICAM-1 governs T-cell exclusion in melanoma"

**Supplementary Table S1. List of antibodies**

| Antibodies | Clone | Manufacturer | Catalog | Application |
|---|---|---|---|---|
| BUV395 anti-mouse CD45 | 30-F11 | BD | 564279 | FC |
| FITC anti-mouse CD45 | 30-F11 | BioLegend | 103108 | |
| BUV737 anti-mouse TCRβ | H57-597 | BD | 612821 | |
| BV605 anti-mouse CD4 | GK1.5 | BioLegend | 100451 | |
| BV650 anti-mouse CD8 | 53-6.7 | BioLegend | 100741 | |
| BV785 anti-mouse NK1.1 | PK136 | BioLegend | 108749 | |
| KB520 anti-mouse F4/80 | RM8 | BioLegend | 123162 | |
| PE anti-mouse CD11b | M1/70 | BioLegend | 101208 | |
| PE-Dazzle 594 anti-mouse Ly-6C | HK1.4 | BioLegend | 128043 | |
| APC Fire750 anti-mouse Ly-6G | 1A8 | BioLegend | 127651 | |
| PE-Cy7 anti-mouse CD19 | 1D3/CD19 | BioLegend | 152417 | |
| AF700 anti-mouse Foxp3 | MF-14 | BioLegend | 126422 | |
| PE-Cy7 anti-mouse CD62L | MEL-14 | BioLegend | 104418 | |
| APC anti-mouse CD44 | IM7 | BioLegend | 103012 | |
| Pacific Blue anti-mouse Granzyme B | GB11 | BioLegend | 612821 | |
| PerCP-Cy5.5 anti-mouse CD11a | M17/4 | BioLegend | 101123 | |
| BUV615 anti-mouse CD31 | MEC13.3 | BD | 752332 | |
| BV605 anti-mouse CD62P | RB40.34 | BD | 740358 | |
| BV786 anti-mouse CD62E | 10E9.6 | BD | 740885 | |
| PerCP-Cy5.5 anti-mouse VCAM-1 | 429 (MVACM.A) | BioLegend | 105715 | |
| APC Fire750 anti-mouse ICAM-1 | YN1/1.7.4 | BioLegend | 116125 | |
| FITC anti-mouse CD31 | MEC13.3 | BD | 01954D | FACS |
| PerCP-Cy5.5 anti-mouse CD45 | 30-F11 | BioLegend | 103131 | |
| anti-mouse CD8α | EPR21769 | Abcam | ab217344 | IF |
| anti-mouse CD31 | 2H8 | Invitrogen | MA3105 | |
| anti-human/mouse Fibrinogen | - | Dako | A0080 | |
| anti-mouse Desmin | - | Sigma | AB907 | |
| Alexa Fluor 488 Donkey anti-Rabbit IgG | - | Jackson ImmunoResearch | 711-545-152 | |
| Alexa Fluor 488 Goat anti-Armenian hamster IgG | - | Jackson ImmunoResearch | 127-545-160 | |
| Alexa Fluor 594 Donkey anti-Rabbit IgG | - | Jackson ImmunoResearch | 711-585-152 | |
| anti-human/mouse CD31 | - | R&D | AF3628 | IHC |
| anti-mouse ICAM-1 | YN1/1.7.4 | BioLegend | 116102 | IHC |
| anti-human CD8 | SP16 | Invitrogen | MA5-14548 | IHC |
| anti-human ICAM-1 | 15.2 | Invitrogen | MA1-80910 | IHC |

FC, Flow cytometry; FACS, Fluorescence-Activated Cell Sorting; IF, Immunofluorescence staining; IHC, Immunohistochemistry

**Supplementary Table S2. Patient cohort characteristics**

| Characteristic | N (%) Total = 37 |
|---|---|
| **Age ranges (median, range)** | 67, (37 - 89) |
| **Gender** | |
| Male | 25 (68%) |
| Female | 12 (32%) |
| **Stage at diagnosis** | |
| Stage 0 | 1 (3%) |
| Stage I | 11 (30%) |
| Stage II | 18 (49%) |
| Stage III | 5 (14%) |
| Stage IV | 1 (3%) |
| **Metastasis occurrence** | |
| Yes | 16 (43%) |
| No | 20 (54%) |
| **Biopsy site** | |
| Primary tumor | 30 (81%) |
| Metastatic tumor | 7 (19%) |
| **Treatment** | |
| ICI | 20 (54%) |
| Non-ICI | 17 (46%) |

Supplementary Figure S1

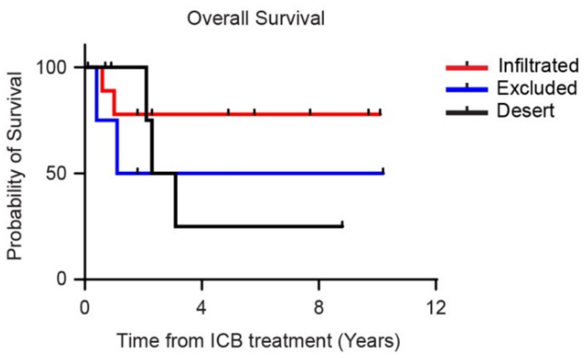


**Supplementary Figure S1. Overall Survival in melanoma patients after ICB treatment.** Kaplan-Meier curves showing overall survival (OS) for patients with three different $CD8^+$ T-cell spatial phenotypes: Infiltrated, Excluded, and Desert. The Desert group showed the worst prognosis, followed by the Excluded and Infiltrated groups.

Supplementary Figure S2

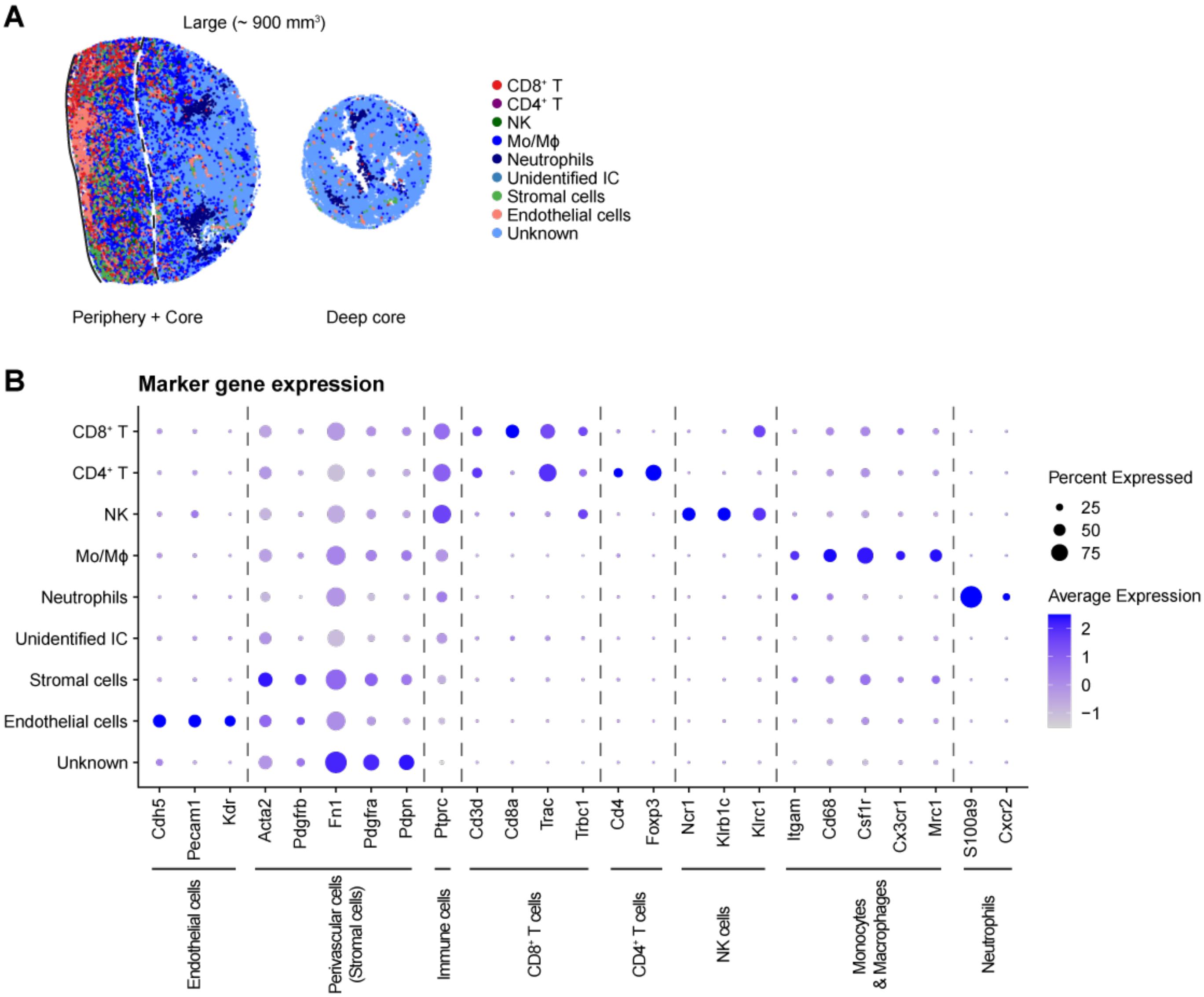


**Supplementary Figure S2. Clustering and marker-based annotation of Xenium spatial transcriptomic data in YUMMER1.7 tumors.** (**A**) Representative Xenium in situ map showing spatially resolved cluster distribution in large YUMMER1.7 tumors (~900 mm$^3$). Distinct immune clusters are enriched at the tumor periphery and excluded from the tumor core and deep-core regions. (**B**) Dot plot showing marker gene expression was used to annotate major immune and stromal cell types. Dot size indicates the proportion of cells expressing each gene, and the color intensity reflects the average expression level.

Supplementary Figure S3

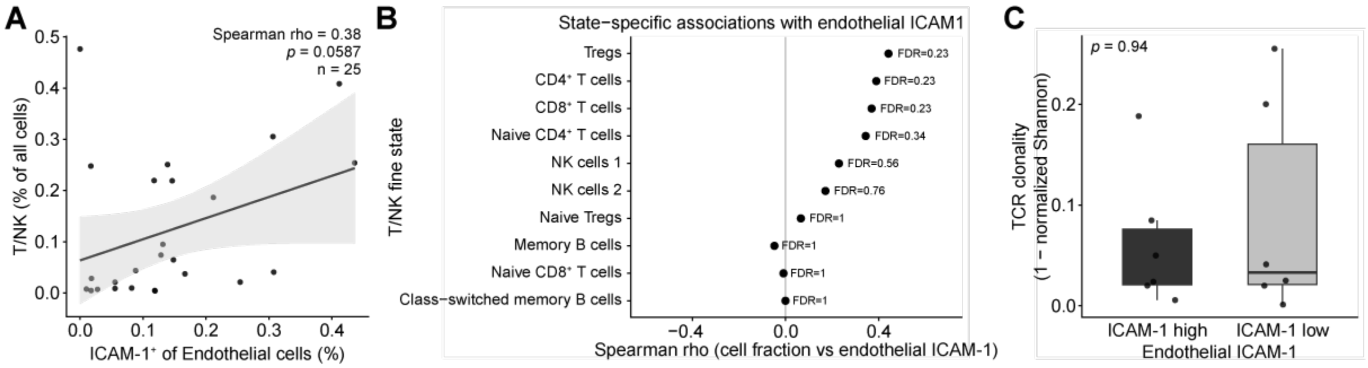


**Supplementary Figure S3. Association of endothelial ICAM-1 with T/NK abundance, T/NK states, and TCR clonality in human melanoma.** (**A**) Scatter plot showing the Spearman correlation between the fraction of ICAM-1$^+$ endothelial cells and total T/NK abundance (fraction of all cells) at the tumor level. Shaded bands represent 95% confidence intervals from linear regression. Each point represents one tumor (n = 25); Spearman's rho (ρ) and two-sided *p*-value are shown. (**B**) Spearman correlations between tumor-level endothelial ICAM-1 expression (mean normalized expression across endothelial cells) and the abundance of annotated T/NK cell states (e.g., CD4$^+$ T, CD8$^+$ T, Treg, NK, and additional subsets). Points represent individual immune states; effect sizes are shown as Spearman ρ; *p*-values were adjusted for multiple testing using the Benjamini-Hochberg false discovery rate (FDR). (**C**) Tumor-level TCR clonality in ICAM-1-high versus ICAM-1-low tumors (upper and lower quartiles of endothelial ICAM-1 expression). Points represent individual tumors; box plots show the median and interquartile range. Statistical comparison was performed using a two-sided Wilcoxon rank-sum test.

Supplementary Figure S4

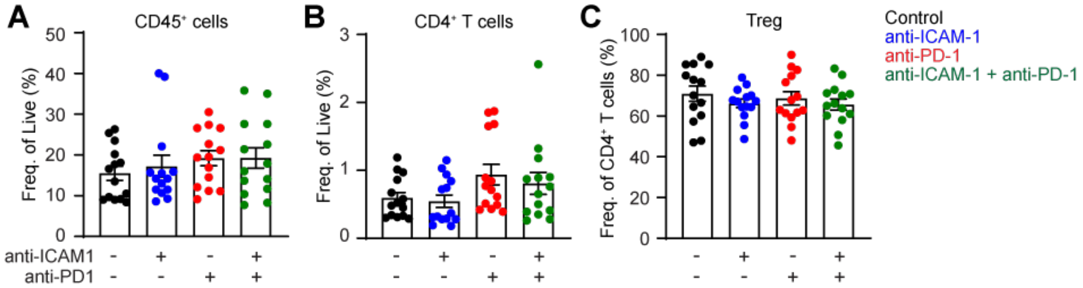


**Supplementary Figure S4. Immune cell infiltration in YUMM1.7 tumors following anti-ICAM-1 and/or anti-PD-1 treatment.** (**A-C**) Flow cytometry analysis of whole tumors showing the frequency of $CD45^+$ immune cells (A) and $CD4^+$ T cells (B) among live cells, and regulatory T cells (Treg; $CD4^+Foxp3^+$) among $CD4^+$ T cells (C). Each point represents one mouse; bars show ± SEM. Statistical analysis was performed using one-way ANOVA.
